# Coordinate-based mapping of tabular data enables fast and scalable queries

**DOI:** 10.1101/536979

**Authors:** Stephen R. Piccolo, Zachary E. Ence, Kimball Hill, PJ Tatlow, Brandon J. Fry, Jonathan B. Dayton

## Abstract

**Motivation:** Biologists commonly store data in tabular form with observations as rows, attributes as columns, and measurements as values. Due to advances in high-throughput technologies, the sizes of tabular datasets are increasing. Some datasets contain millions of rows or columns. To work effectively with such data, researchers must be able to efficiently extract subsets of the data (using filters to select specific rows and retrieving specific columns). However, existing methodologies for querying tabular data do not scale adequately to large datasets or require specialized tools for processing. We sought a methodology that would overcome these challenges and that could be applied to an existing, text-based format.

**Results:** In a systematic benchmark, we tested 10 techniques for querying simulated, tabular datasets. These techniques included a delimiter-splitting method, the Python *pandas* module, regular expressions, object serialization, the *awk* utility, and string-based indexing. We found that storing the data in fixed-width formats provided excellent performance for extracting data subsets. Because columns have the same width on every row, we could pre-calculate column and row coordinates and quickly extract relevant data from the files. Memory mapping led to additional performance gains. A limitation of fixed-width files is the increased storage requirement of buffer characters. Compression algorithms help to mitigate this limitation at a cost of reduced query speeds. Lastly, we used this methodology to transpose tabular files that were hundreds of gigabytes in size, without creating temporary files. We propose coordinate-based, fixed-width storage as a fast, scalable methodology for querying tabular biological data.

## Introduction

Biologists often generate data suitable for representation in an attribute-value system^1^, also known as an information system^2^, simple frame^3^, object-predicate table^4^, or flat file. In this representation, an *object* might be a biological organism, an *attribute* might be a characteristic of that organism, and a *value* might be a datum for that object and attribute. For example, a researcher might observe 200 cancer patients (objects) and collect transcriptomic measurements for 20,000 genes (attributes); each *value* would indicate the relative number of transcripts present in tumor cells for each patient/gene combination^5^. In this example, the data values would have been summarized previously using preprocessing tools, such as a reference aligner and a transcript-quantification algorithm^6–9^. For convenience and compactness, researchers typically store attribute-value data in 2-dimensional, tabular formats. Commonly, in such tables, each row contains data for a given object, and each column contains data for a given attribute^10^; but in some cases, the table is transposed (objects as columns, attributes as rows). Researchers use tabular data to perform analytical tasks, such as executing statistical analyses, producing graphics, and further summarizing the data.

Across the subfields of biology, researchers store a considerable proportion of tabular data in plain-text formats. This approach coincides with the Unix and “Pragmatic Programming” philosophies^11–13^, which advocate for storing data and sharing data among computer programs as plain text. Keeping data as text has many advantages. Plain text is readable to humans.Sophisticated text editors are freely available for all major operating systems. A wide range of tools exist for generating, manipulating, parsing, and compressing text files; these include long-established command-line tools developed by the Unix community. In addition, scripting languages like Python^14^ and R^15^ provide libraries for analyzing text-based tabular data; these libraries are used broadly within the biology community and elsewhere^16^. Storing data as plain text does have drawbacks relative to binary formats. Text files may be larger than binary files, and it may be more computationally intensive to parse a text file than a binary file. Consequently, a multitude of techniques for storing tabular data in binary (non-text) formats has been developed. For example, researchers use Microsoft Excel for data exploration and analysis^17,18;^ relational databases provide a formalized methodology to query tabular data^19^; so-called NoSQL databases provide alternative methodologies for structuring and querying data, including attribute-value systems^20^; the Hierarchical Data Format (HDF5) is often used for tabular data, including in biology research^21,22^. Additionally, in recent years, distributed architectures for large-scale data storage and processing have seen wide use; these technologies include Apache Hadoop and Apache Spark^23,24^. Despite these advances, the humble plain-text file continues to play a critical role in biology research due to its simplicity, flexibility, familiarity, and portability.

In our own research studying molecular profiles of tumors—and via collaborations with other scientists—we have frequently encountered a need to *select* and *project* tabular data. In relational-algebra terms^19^, selection refers to the process of identifying rows that match some criteria; projection refers to the process of retrieving specific columns. For example, in a study of genomic and transcriptomic profiles of human breast tumors, a researcher might wish to select only patients diagnosed before the age of 40 and who harbor a mutation in *BRCA1*, a gene known to effect double-stranded DNA repair by homologous recombination^25^. Having identified this patient subset, the researcher might wish to retrieve (project) transcriptomic data for genes that interact with *BRCA1*. In repositories like The Cancer Genome Atlas (TCGA), The International Genome Consortium (ICGC), and Gene Expression Omnibus (GEO)^26–28^, genomic and transcriptomic data—and their corresponding annotations—are stored in tabular text files. The ways that objects, attributes, and values are oriented within these files differ across these and other repositories^29–31^, but values are commonly oriented in rows and columns and are separated by tab characters, comma characters, or some other delimiter.

Typically, to parse such data, researchers write custom scripts or use software packages that facilitate parsing^32–34^. To perform selection, the code must extract all values from the column(s) to be used as filtering criteria. If data values are delimited by tab characters, for example, the code must identify the positions of tab characters and extract data at the relevant positions for each row. However, because data values may vary in length, the positions of tab characters may differ for each row, and these positions must be reidentified for each row, thus slowing execution. After identifying rows that match the selection criteria, the researcher may then wish to project the data. When parsing a tab-delimited file, the code must again identify positions of tab characters *for each row* and extract values at the relevant positions. In this methodology, the code parses the data row by row, thus minimizing memory consumption. Alternatively, the entire file could be parsed into an in-memory data structure; this methodology may increase the efficiency of selection and projection, but many datasets are too large to fit in memory. Additionally, if a researcher wishes to use only a few columns for selection or projection, it is inefficient to read the entire file into memory. Hybrid solutions exist, such as the *pandas* module for Python^35^. However, in our experience, it has been difficult to find a solution that strikes a satisfactory balance between speed and memory usage. This challenge has become more acute as data sizes have increased. For example, a re-quantification of RNA expression data from TCGA contains data for 11,373 tumors across 199,169 transcripts^36^.Phase I of the Library of Integrated Network-based Cellular Signatures (LINCS) yielded data for more than 1.3 million experiments, including transcriptomic data for 12,300 genes (after imputation) and annotations for each experiment^37^. Recently, the UK Biobank posted genotypic, phenotypic, and health-related measurements for approximately 500,000 individuals^38^. Some of these files are multiple terabytes in size. These trends are true in other fields as well, including proteomics, remote sensing, and imaging^39–42^.

In evaluating methodologies that could handle such data, we envisioned scenarios in which data files are created once and then queried many times. Public repositories like TCGA, LINCS, and UK Biobank cater to these scenarios; after the data have been prepared, they are stored on web servers, enabling researchers to download and query the data. Because the files are written only once, it is less important to optimize speeds for writing the files, and it is unnecessary to support concurrent writing by multiple agents. In contrast, it is highly preferable that researchers can query the data quickly and flexibly. With this context in mind, we sought a solution that would meet the following criteria:

- Handle datasets larger than what can fit into memory on modern personal computers.
- Handle attributes of different types (categorical, ordinal, numeric, etc.).
- Support selection based on data in *any* column.
- Store the data in a portable format that can be transported across systems without custom tools or specialized expertise.
- Store the data in a space-efficient manner (while preferring fast speeds over reduced storage).
- Not be specific to any particular type of biological data (e.g., genomic, transcriptomic, ecological).
- Can be created, indexed, and queried in a non-proprietary^43^, programming-language agnostic, and platform-independent manner.
- Can transpose rows and columns without reading *all* the data into memory and without creating temporary files.
- Can represent missing values explicitly.

In our quest to identify a solution that would address these criteria, we considered a variety of binary-based solutions. These included relational databases, NoSQL databases, HDF5, and the Apache Parquet format^44^. With each solution, we faced limitations. For example, the SQLite relational database has a limit of 32,767 columns^45^. NoSQL databases provide many options for structuring the data, but we failed to identify an approach that would provide adequate query speeds and storage sizes. The HDF5 format is designed primarily for numerical data, whereas we sought the ability to handle other data types as well. As a columnar storage solution, Parquet was efficient at projection; however, it was ill-suited to selection. Ultimately, we focused on text-based solutions, performing a benchmark analysis of 10 different techniques for parsing tabular data. As described below, we chose one of these techniques and refined it further. We found that this technique addresses each of the above criteria yet is human readable, fast, and scalable.

## Methods

In an initial round of benchmarks, we used Python scripts to generate tabular text files in which 10% of the columns contained categorical values (randomly generated, 2-digit alphabetical sequences) and 90% of the columns contained numerical values (ranging between 0.0 and 1.0). First, we used these scripts to generate relatively small files, containing 100 columns and 1000 rows. After verifying functionality, we generated two types of large file that represent dimensions that will be increasingly seen in biological research: 1) “tall” files containing 1 million rows and 1,000 columns, and 2) “wide” files containing 1,000 rows and 1 million columns. Each of these files contained a total of 1 billion data points (approximately 10 GB in size). For each set of dimensions, we saved the data in four different formats:

- *tsv*. We separated each value on each row with tabs (tab-separated-value format).
- *msgpack*. We used the MessagePack format^46^ to serialize each row of data as a list object.
- *flags*. In an attempt to make it faster to access elements at a given column index, we specified the index of each element within each row of data and embedded these indices within the file, prior to each datum.
- *fwf*. The width of each column corresponded to the data value with the largest number of characters in that column (fixed-width format). We also added a buffer character between columns.

In this phase, we evaluated 10 techniques for projecting the data. Different techniques used different versions of the input data (see below). We coded each technique to select the first column and every hundredth column thereafter. Each script saved the selected columns to a tab-delimited text file. We then used a script to verify that the output was correct.

- *delimiter-split*. We used TSV files as input, split each line on tabs, and extracted values at the specified indices.
- *pandas*. We applied the *read_csv* function from the Python *pandas* module to the TSV files.
- *reg-ex-quant*. We used regular expressions to quantify tab characters that preceeded each specified index and then extracted those values using capturing groups.
- *reg-ex-tab*. We used regular expressions to map non-capturing groups to indices that should be ignored and capturing groups to indices that should be extracted.
- *msgpack*. We deserialized each row from the MessagePack serialized files and extracted values at the specified indices.
- *flags*. For the flag files, we identified the position of each specified flag and then extracted characters after it until another flag was reached.
- *awk*. We applied the *awk* command-line utility to the TSV files^47^. This Unix-based tool provides extensive support for parsing text files.
- *gawk*. This is another variation on awk.
- *nawk*. This is yet another variation on awk.
- *fixed-width*. We used the header line in the file to identify the starting and ending positions of each column and then used string indexing to extract values at the specified indices.

Aside from the *awk*-based solutions, we used Python code. In addition, for each Python solution, we implemented a memory-mapping version of the code. Memory mapping supports the ability to randomly access locations within a file; accordingly, in some cases, we could extract specific portions of the file without needing to iterate sequentially through the file or read every character into memory.

During the second phase of this study, we developed a modified version of the fixed-width format (*fwf2*). First, we calculated the position and width of each column and stored these values in an index file that we could also memory map. Second, we calculated the full length of the first line—all lines should have the same length—and stored this length in a second index file. These changes enabled us to quickly calculate row and column coordinates when querying the data. Lastly, we removed the extraneous buffer characters between the columns. This reduced file sizes; however, we retained a nonessential newline character at the end of each line to make the files more readable.

All of our code, along with a bash script to execute the benchmarks, can be found at https://github.com/srp33/Tabular_File_Benchmark. The same repository contains an R Markdown file that includes the code we used to create figures for this paper. We used R version 3.5.1 and the *ggplot2, readr, dplyr*, and *cowplot* packages for the figures^48–51^. For the benchmarks, we used Python (version 3.6.7) and the following external Python modules: msgpack (0.5.6), numpy (1.15.2), pandas (0.23.4), and snappy (0.5.3). All benchmark tests were executed on a 64-bit processor running Ubuntu Linux (18.04) with the following hardware specifications:

- 4.5 GHz Xeon^®^ W-2155 (3.3 up to 4.5 GHz – 10 Cores - 20 Threads - 2666 MHz)
- 256 GB Quad Channel DDR4 random access memory at 2666 MHz (8× 32GB)
- 250 GB NVMe PCIe M.2 solid-state drive (SSD) for the operating system
- 3.8 TB NVMe 3.84TB U.2 Mixed Use SSD for data storage

## Results

First, we evaluated methods for projecting tabular text files that contained 1 billion data points. These files had either a “tall” or “wide” orientation. The tall files simulate scenarios in which researchers collect 1,000 data points for 1 million patients (or other object type). The wide files simulate scenarios in which researchers collect 1 million data points for 1,000 patients. As high-throughput data-generation technologies advance and as researchers combine individual datasets into aggregate ones, such scenarios will be increasingly common.

Commonly, biology researchers store data in tab- or comma-delimited files and parse such files using the *delimiter-split* method. Thus, we considered the performance of this approach to be a baseline. For the tall files, our scripts extracted every hundredth column in 21.72 and 17.76 seconds with and without memory mapping, respectively. All but two of the competing methods outperformed this approach (Figure 1). In contrast, on wide files, the performance slowed considerably for all methods. The baseline method extracted every hundredth column in 31.38 and 27.44 seconds. The *pandas* method and both regular-expression methods performed worse than the baseline, and their performance was dramatically worse than it was on the tall files. The poor performance of *pandas* is perhaps surprising, given the package’s popularity among data scientists^52^.

**Figure 1:**
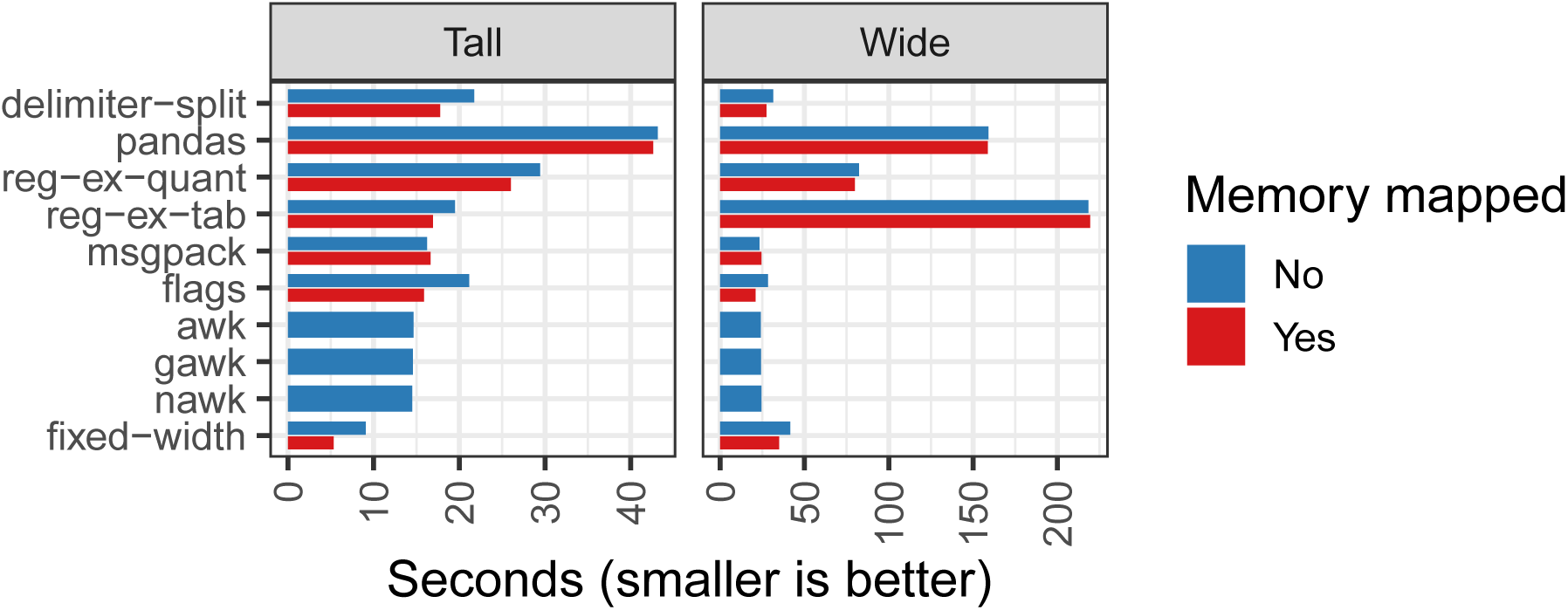
Execution speed for 10 methods of parsing tabular text files. We evaluated techniques for projecting every hundredth column from tabular text files. The techniques varied in the ways that text files were structured and how they were parsed (see Methods). *awk, gawk*, and *nawk* did not support memory mapping. Tall files consisted of 1 million rows and 1,000 columns; “Wide” files had 1,000 rows and 1 million columns. Each file included a mixture of categorical (10%) and numerical (90%) attributes. Note that the x-axis scales differ for the two panels.

The fixed-width method performed best overall on the tall file, projecting the data in only 5.34 seconds with memory mapping; however, its performance was mediocre on the wide file. We hypothesized that a few adjustments to the file format and our algorithmic approach might improve the performance substantially (see Methods). In addition, we implemented a chunking scheme in which we parsed 1000 rows of input data at a time before writing to the output file. After these adjustments, we projected every hundredth row from the tall file in 3.70 seconds. For the wide file, we projected the data in 3.43 seconds, only 10% the duration of the original approach. Given these results, we focused on this method and evaluated its performance further.

We wanted a method that would excel at projection *and* selection. Therefore, we performed selection on data from one column with categorical data and one column with numerical data. The categorical values were 2-character sequences of letters; arbitrarily, we searched for values that started with “A” or ended with “Z”. The numerical filter searched the specified column in the remaining rows for values greater than or equal to 0.1. These criteria yielded approximately 6.9% of the rows. Lastly, we projected every hundredth column. This process took 1.04 seconds for the tall file and 0.28 seconds for the wide file.

When performing the initial benchmarks, we stored the data in four tabular formats (see Methods). The *flags* and *fixed-width* files were larger than the other formats, especially for the wide files (see Figure 2). To enable indexing, these formats require extra text within the files. We considered ways to reduce this extra storage requirement while still supporting fast query times. We tested four compression algorithms: *gzip, bzip2, lzma*, and *snappy*. We compressed the text files line by line. After compression, the lengths of the lines varied, so we saved the starting position of each row to a serialized dictionary. Compression times differed considerably across the methods (Figure 3); as its name implies, *snappy* was extremely fast. In constrast, *snappy*-compressed files were approximately twice as large as files compressed using the other algorithms. Most importantly for this study, select-and-project speeds were dramatically faster for *snappy*-compressed files than for any of the other algorithms (Figure 3). However, these speeds are 20-50 times slower than we attained using non-compressed data. We needed to decompress each full line before we could evaluate the selection criteria or perform projection. Accordingly, individuals who consider using this methodology must consider the substantial tradeoff between speed and space requirements.

**Figure 2:**
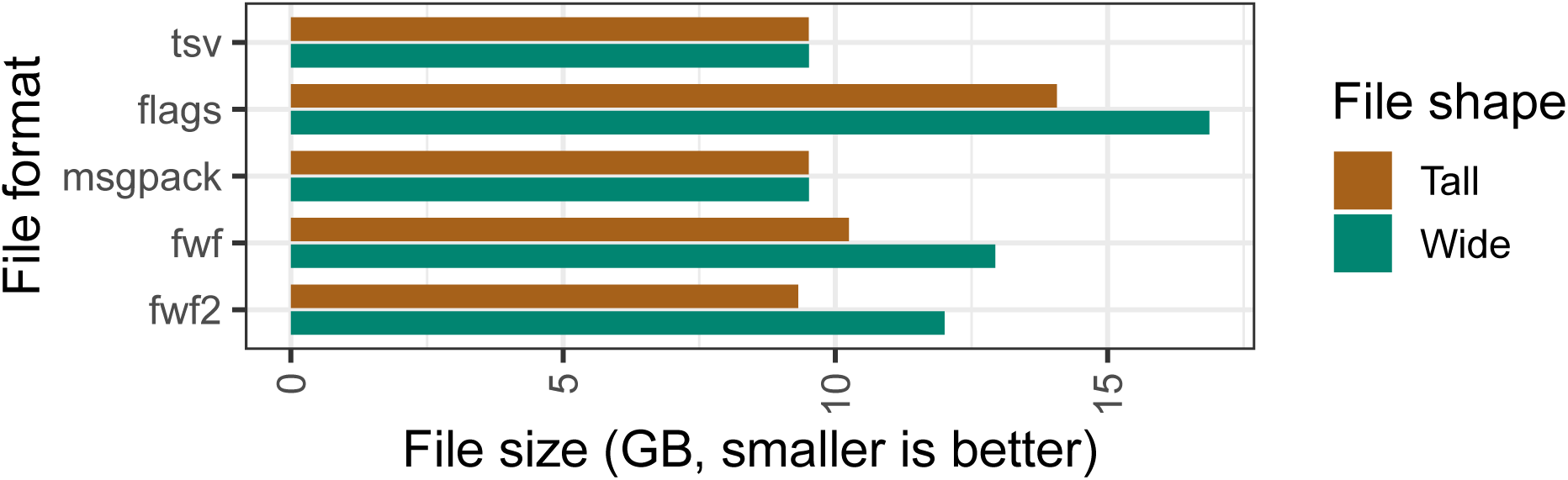
Sizes of simulated data files used in the initial benchmarks. *tsv* = Tab-separated values. *Flags* = A “flag” before each value indicates the column index of each value. *msgpack* = Each row was serialized as a Python list into MessagePack format. *fwf* = Fixed-width format. *fwf2* = Modified fixed-width format.

**Figure 3:**
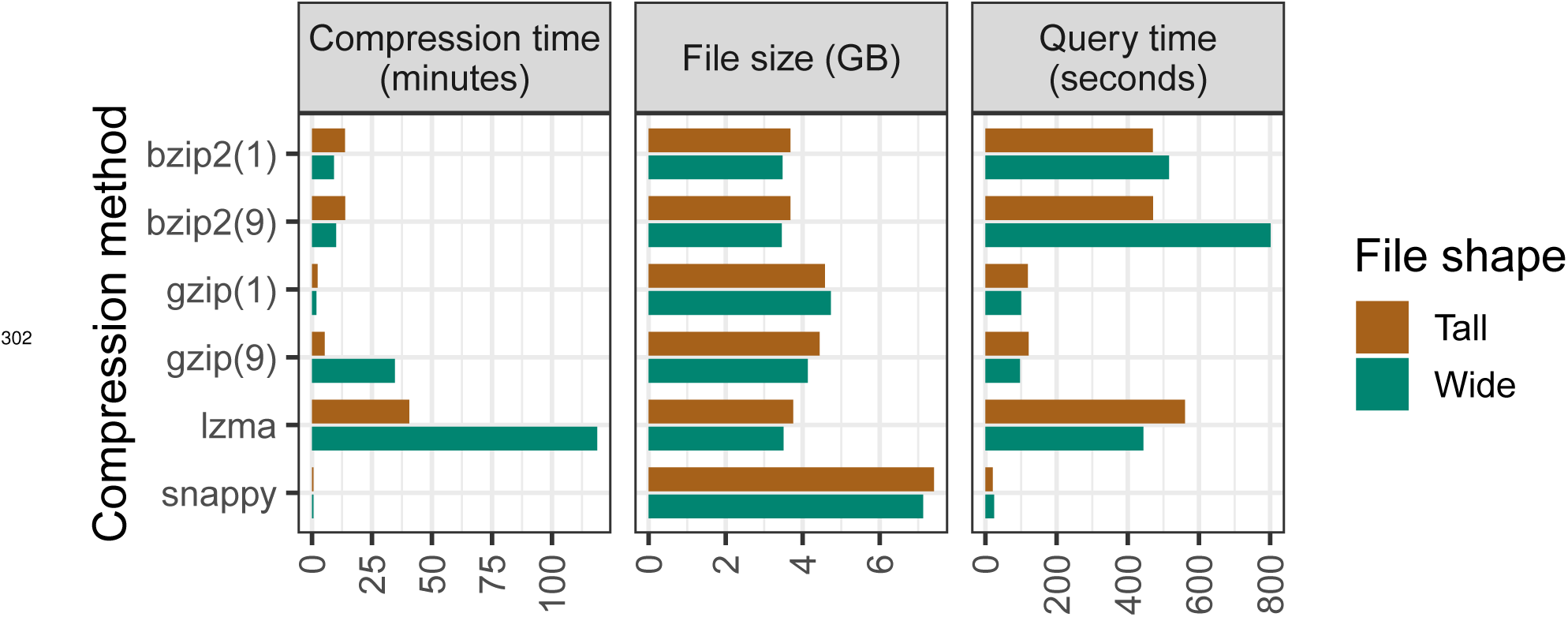
Results of compression benchmarks. In an attempt to reduce file sizes, we compressed the fixed-width (fwf2) files line by line and stored the starting position of each line to enable faster file traversal. The first panel shows how long it took to compress the files. The second panel indicates file sizes after compression (the original files were approximately 10 GB in size). The third panel illustrates how long it took to select and project data from the compressed files. The gzip and bzip2 algorithms support a parameter to alter the level of compression; we used levels 1 and 9, which are indicated in parentheses. Although the *snappy* compression algorithm was much faster than the other algorithms, these speeds were 20-50X slower than without compression.

Next we tested the scalability of the (non-compressed) fixed-width approach. We simulated a scenario in which a researcher might wish to store genotypes for a large number of individuals. We generated text files that contained simulated genotypes with 10, 50, 100, 500, 1000, 5000, 10000, 50000, 100000 or 500000 rows and columns, respectively. In these files, we represented genetic loci as rows and organisms as columns and used a pair of nucleotide characters (A, C, G, T) to represent each genotype. We tested our ability to select and project the data by extracting genotypes for the intersection of 10 random rows and 10 random columns. Query speeds were identical (0.03 seconds) for all file sizes except the largest (0.06 seconds). The largest file (500000 rows by 500000 columns) had a total of 250 billion data points and was 465 GB in size (Figure 4A). Although extracting 10 rows and 10 columns does not reflect a real-world scenario, it illustrates the promise of performing these operations quickly on extremely large files.

**Figure 4:**
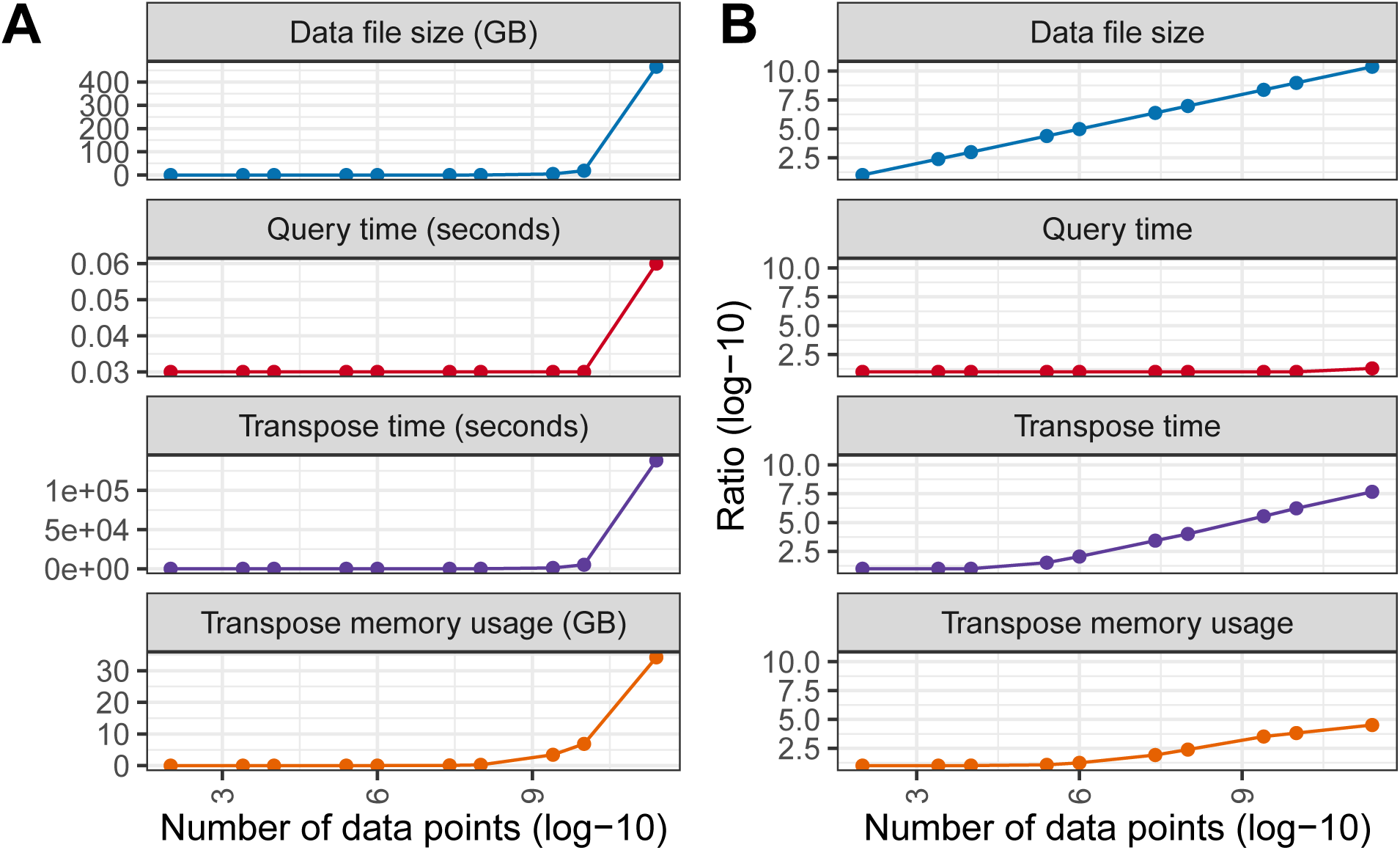
Results of simulated genotype benchmarks. We simulated genotypes for cohorts and genomes of increasing size. The x-axes indicate the total number of simulated genotypes. In **A**, the y-axes indicate *absolute* performance for each metric. In **B**, the y-axes indicate performance *relative* to what was observed for the minimum number of genotypes. File size increased linearly, whereas the other metrics, especially query time, increased at a slower rate.

As a final test, we transposed the simulated genotype files. To our knowledge, no tool exists for transposing tabular files without reading the full dataset into memory or writing temporary files. Our fixed-width approach successfully transposed each of the simulated genotype files, including the largest, without writing temporary files. Time and memory usage did increase as file sizes increased, but in a nonlinear fashion (Figure 4B).

## Discussion

Our goal was to evaluate techniques for storing and querying tabular data. Such data are used widely in biology research. We sought to identify a methodology that would reduce query times and overcome limitations of existing approaches. In addition, we sought a solution that would provide the flexibility and readability of plain text plus some benefits of binary formats. Our results suggest that fixed-width formatting, memory mapping, and coordinate-based indices can meet these needs for many research scenarios, especially those that prioritize fast reading over fast writing and that can accept larger file sizes as a compromise for faster querying. In this study, we simulated data for which all values in a given column are the same width; but in practice, data values in a given column often vary in length. In these scenarios, extra buffer characters are needed. More research is necessary to evaluate how much these buffer characters would increase file sizes in practice, but we predict that query speeds will be impacted only minimally. One possibility for mitigating the effects of larger file sizes is to compress the whole file using a standard compression scheme (e.g., gzip) before it is placed on a web server; accordingly, distributing the file would require less disk space on the server and less network bandwidth during file transfer; the researcher could then decompress the file locally before querying it.

The results of time-based benchmarks must be interpreted with caution because execution times vary from one computer system to another. Additionally, we performed these benchmarks on hardware whose performance exceeds that of many computer systems used currently for biology research. However, we are confident in our conclusions about the *relative* performances of the methods we evaluated.

This study describes a proof of concept rather than a production-ready tool. One limitation of our current approach is that data types are not stored explicitly for each attribute; these must be inferred from the data. However, we believe our methodology’s performance in these benchmarks merits further development.

The simplicity of our methodology is one of its strong points. It should be possible to implement this approach in any programming language and operating system that support reading and writing text files as well as memory mapping. We do not intend this to be “yet another file format” for bioinformaticians to deal with; rather, we describe it as a methodology for extending an existing format. In addition, our approach could facilitate translation among formats and data orientations. Further methodological refinements are possible, potentially including more sophisticated algorithms for identifying column widths and compressing the data. We invite collaborations with others in the research community.

